# Peripheral direct current suppresses physiologically evoked nociceptive responses at the spinal cord in rodent models of pain

**DOI:** 10.1101/2023.06.07.544048

**Authors:** Tom F. Su, Jack D. Hamilton, Yiru Guo, Jason R. Potas, Mohit N. Shivdasani, Gila Moalem-Taylor, Gene Y. Fridman, Felix P. Aplin

## Abstract

Electrical neuromodulation is an established non-pharmacological treatment for chronic pain. However, existing devices using pulsatile stimulation are not suitable for all types of chronic pain. Direct current stimulation is a recently developed technology which shows better selectivity for small-diameter fibres. We investigated if this selectivity could be applied to preferentially suppress nociceptive signalling. We applied direct current to the sciatic nerve of rats and compared its effects on spinal activity produced by physiological (non-electrical) stimuli delivered to the foot. Tests were performed across models of neuropathic and inflammatory pain to further clarify potential clinical applications. We found that direct current could effectively suppress activity relating to painful stimuli in both pain models tested. These findings strongly support the use of direct current neuromodulation for chronic pain relief, and suggest that it may be effective at treating a broader range of aberrant pain conditions than existing devices.

## Introduction

Chronic pain, a debilitating condition affecting roughly one in five people (1), is defined as pain that persists for at least 3 months (2). Chronic nociceptive pain is caused by damage or inflammation in non-nervous tissue, while chronic neuropathic pain is caused by lesions within the somatosensory system itself (3). First-order drugs for both subtypes of chronic pain are not well suited to chronic prescription due to drawbacks such as high risk of dependence in opioids (4, 5) or loss of efficacy over time in gabapentinoids (6). Electrical neuromodulation devices such as spinal cord or peripheral nerve stimulators offer a non-pharmacological alternative for chronic pain management (7–10). Although many such devices have received approval for clinical use, they only provide relief for a subset of patients (11, 12), and there is limited evidence to support their efficacy for chronic nociceptive pain specifically (13, 14). A potential explanation for these shortcomings lies in the waveforms used for neuromodulation; existing stimulators deliver short electrical pulses which primarily affect large-diameter, primarily non-nociceptive Aα/Aβ fibres and leverage spinal gating mechanisms to indirectly modulate nociceptive signalling (15, 16). An alternative neuromodulation paradigm which instead targets the activity of small-diameter, nociceptive Aδ/C fibres could therefore allow devices to provide pain relief in a wider range of contexts without interfering with touch sensations necessary for everyday life.

Direct current (DC) neuromodulation may provide this alternative, as there is a growing body of evidence to suggest that small-diameter neurons are sensitive to DC and other low frequency waveforms (17, 18). DC has historically been avoided in clinical devices due to cytotoxicity concerns (19, 20), but recent engineering developments have paved the way for safe in vivo DC delivery (21–23). Indeed, neuropathic pain relief using ultra-low frequency dorsal column stimulation has been demonstrated in a recent clinical trial (24). A different approach using peripheral DC wherein nociceptive signalling is selectively disrupted before reaching the spinal cord has also been proposed (18, 23, 25). This approach could be particularly beneficial for the treatment of chronic nociceptive pain, such as post-surgical or arthritic pain, which is not well managed by existing neuromodulation devices (13, 14).

However, it is unknown if peripheral suppression of small-diameter fibre activity translates to targeted suppression of pain pathways. Previous studies characterised fibre types by electrically evoked response latency, rather than by sensory input (18, 24). While there is a correlation between response latency and function, this relationship is disrupted in chronic pain conditions where large-diameter A fibres contribute to pain hypersensitivity via mechanical allodynia and hyperalgesia (26, 27). Clarifying the functional, rather than anatomical, populations targeted by DC suppression is important for evaluating the therapeutic potential of peripheral DC neuromodulation. Furthermore, given that chronic nociceptive and chronic neuropathic pain promote different pathological changes within the nervous system (28), it is necessary to compare the effects of DC neuromodulation across both pain conditions.

In this paper, we compare the effects of peripheral DC neuromodulation in rodent models of neuropathic and inflammatory nociceptive pain by recording spinal responses to functionally relevant peripheral stimuli. Based on previous findings which measured electrically evoked activity (18), we hypothesised that responses evoked by noxious stimuli would be preferentially suppressed in all treatment groups. We found that noxious-evoked responses and potentially allodynic tactile-evoked responses were suppressed preferentially in both pain models, but not in naïve animals. Furthermore, we demonstrated that conduction block persisted after application of DC in nociceptive responses only. Our findings support the continued translation of peripheral DC neuromodulation devices as a potential treatment for both nociceptive and neuropathic pain.

## Results

We first verified the presence of mechanical allodynia and thermal hyperalgesia at the hindpaw in the neuropathic (spared nerve injury) and inflammatory (complete Freund’s adjuvant) pain models using behavioural testing. Figure 1A shows painful mechanical force threshold (von Frey) and thermal withdrawal latency (Hargreaves) for both hindpaws pre- and post-treatment. In the ipsilateral (treated) hindpaw, we observed a significant increase from baseline in mechanical sensitivity for both models (p < .0001); thermal sensitivity increased significantly for the inflammatory pain model only (p < .0001). There were no significant changes in sensitivity on the contralateral (untreated) side.

**Figure 1:**
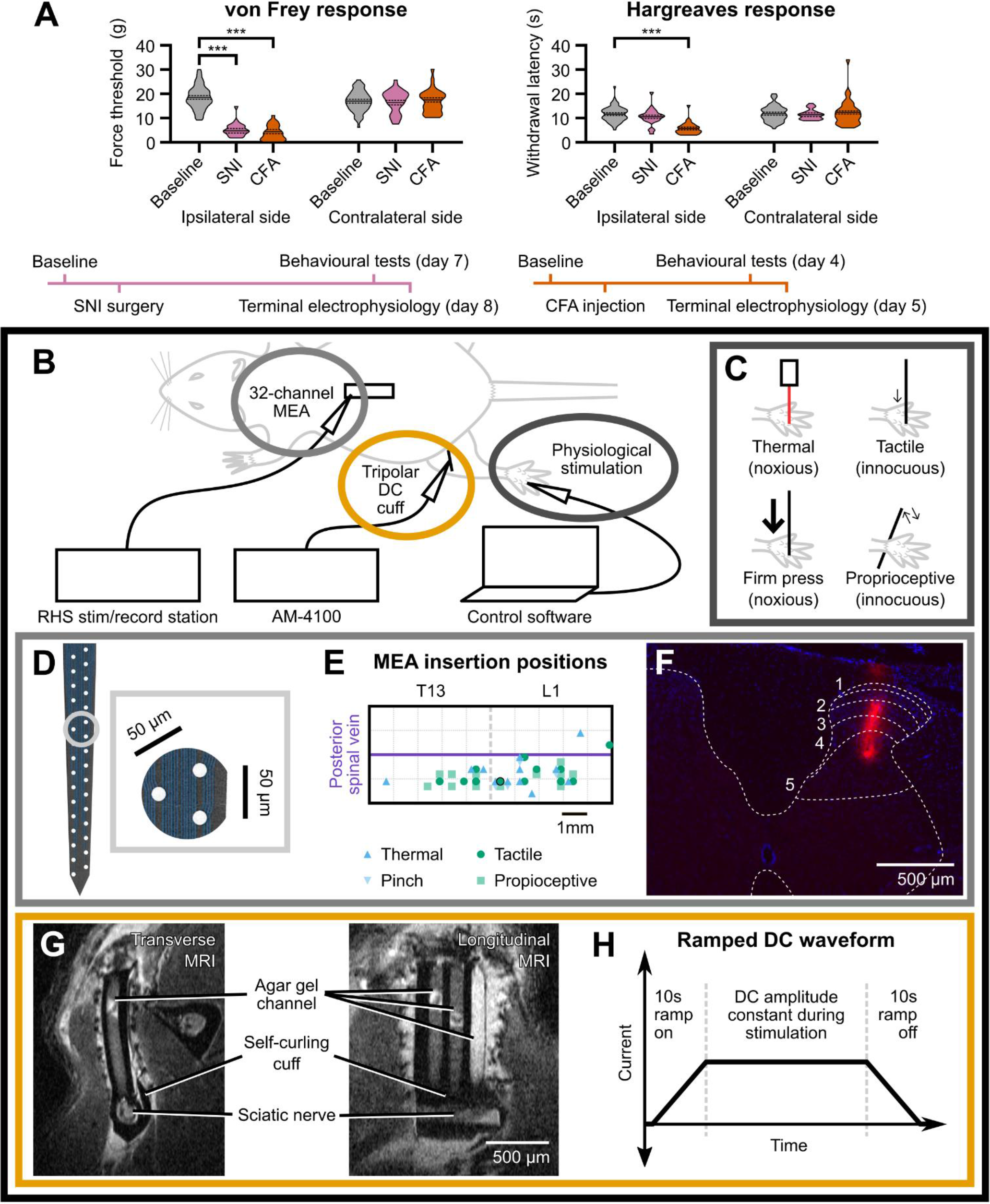
Experimental design. (A) Behavioural verification of pain models. von Frey force thresholds and Hargreaves response latencies for both hindpaws in each treated animal, at baseline and one day before terminal experiment. Time courses for each pain model are depicted below plots. Force thresholds for the ipsilateral hindpaw were significantly lower than baseline in all treated animals, while response latencies were lower in the ipsilateral hindpaw of CFA-treated animals only. EMM ± SE is shown. ***: p < .001. (B) Electrophysiological setup. Spinal recordings were taken using a 32-channel MEA and an RHS Stim/Record System. DC was delivered to the sciatic nerve by an AM-Systems stimulator via a tripolar cuff. The hindpaw was stimulated physiologically with the aid of software triggers. (C) Physiological stimulus types. Four physiological stimuli were chosen to evoke responses that were noxious thermal–, noxious press–, tactile-, or proprioceptive-dominant. (D) MEA layout. NeuroNexus A1×32-Poly2-5mm-50s-177 MEA with 50 μm electrode spacing. Inset: example three-channel set chosen for spike-sorting. (E) MEA insertion positions. MEA was inserted into T13/L1 spinal segments ipsilateral to the stimulation site. (F) DiI stain of MEA insertion highlighted in black. Red fluorescence shows depth and medio-lateral position. Dorsal horn and approximate L1 spinal laminae positions are marked. (G) Transverse and longitudinal MRI scans of implanted tripolar cuff. (H) Ramped DC waveform delivered via tripolar cuff.

Having confirmed the development of pain hypersensitivity, we collected physiologically evoked spinal recordings (Figure 1B). We evoked responses at the hindpaw by applying four physiological stimulus paradigms (noxious thermal–, noxious press–, tactile-, and proprioceptive-dominant; Figure 1C). These responses were recorded at T13-L1 using a 32-channel multi-electrode array Figure 1D-E). Cross-sectional DiI fluorescence confirmed that the electrode array penetrated laminae 1-5 of the dorsal horn (Figure 1F). To test the effects of DC neuromodulation on these responses, we implanted a tripolar silicone cuff around the ipsilateral sciatic nerve through which we delivered ramped DC waveforms. Figure 1G shows transverse and longitudinal MRI scans of an implanted cuff. The DC waveform delivered is depicted in Figure 1H.

We visualised recorded spike activity and selected channels for further analysis based on the presence and volume of post-stimulus spike activity. Figure 2 shows representative voltage traces in naïve animals corresponding to each of the four stimulus paradigms. Each stimulus paradigm evoked a distinctly identifiable response pattern. Noxious thermal–dominant responses (Figure 2A) were characterised by sustained firing and a latency of ∼1 second from stimulus onset due to the extended time course of the stimulus itself. Noxious press–dominant responses (Figure 2B) were characterised by wide but well-defined bursts of activity, with a latency of 50-100 ms from stimulus onset. Tactile-dominant responses (Figure 2C) were typically single-or double-spike events, with highly consistent timings within 10ms of the stimulus. Proprioceptive-dominant responses (Figure 2D) followed a bimodal burst pattern caused by rotation and counter-rotation of the joint across 400 ms. We then spike-sorted these responses into units using principal components analysis across three channels. This yielded 523 units from naïve animals (n = 27 animals), 292 units from the neuropathic pain group (n = 8 animals), and 300 units from the inflammatory pain group (n = 12 animals).

**Figure 2:**
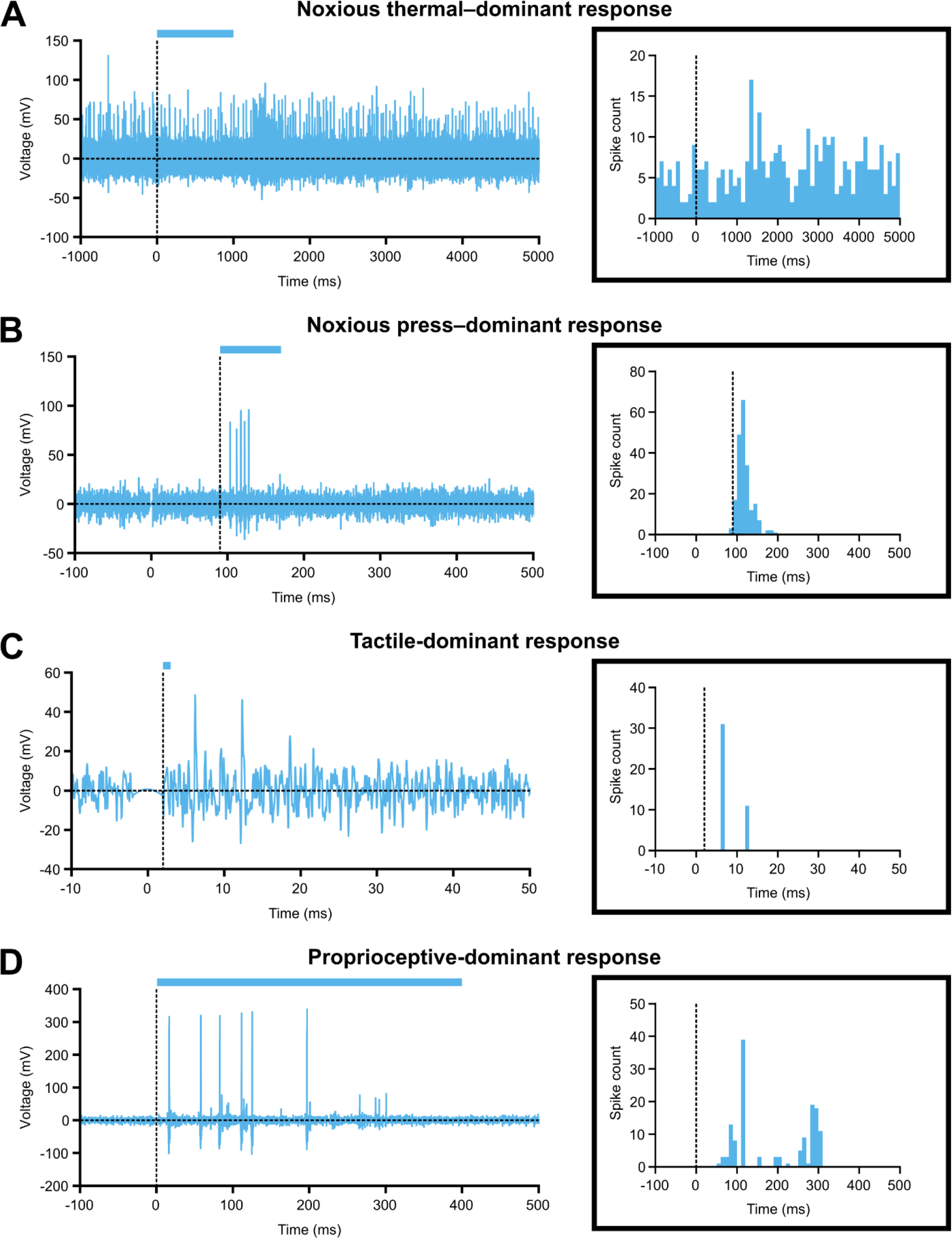
Example signal traces of each physiological stimulus type. Representative dorsal horn responses to (A) noxious thermal–, (B) noxious press–, (C) tactile-, and (D) proprioceptive-dominant stimuli. Vertical dotted line indicates start of stimulus, and horizontal coloured bar represents duration of stimulus. Examples are taken from naïve animals. Inset: histograms of single units obtained from the corresponding trace by spike-sorting. Histogram bins are 1/60th of the displayed time (i.e. 100ms for noxious thermal–dominant response, 10ms for noxious press– and proprioceptive-dominant responses, and 1ms for tactile-dominant response). Note that Y-axis is different for each plot to improve readability.

### Peripheral DC application suppresses spinal cord responses evoked by pain-related stimuli in rodent models of pain

To determine the effects of sciatic nerve DC neuromodulation on sensory inputs to the spinal cord, we examined the activity of spike-sorted units before and after application of DC. Figure 3 shows spike count histograms of representative units during 1000 μA DC (yellow) overlaid on their pre-DC baseline activity (blue). Rasters of the same units can be found in Supplementary Materials (Figure S1). Each row corresponds to a physiological stimulus paradigm, while each column corresponds to a treatment group. Note that to better highlight DC-mediated changes in response pattern, bin widths are consistent across each row but not between rows, and y-axis scale varies between each histogram. A key observation shown in this figure is that many tactile-dominant responses in pain models did not exhibit the characteristic response pattern of naïve tactile-dominant units, displaying instead a longer peak response latency (Kruskal-Wallis; p < .001 for neuropathic pain group, p = .046 for inflammatory pain group) and a non-unimodal latency distribution as indicated by Hartigans’ dip test (Table 1; 29, 30).

**Figure 3:**
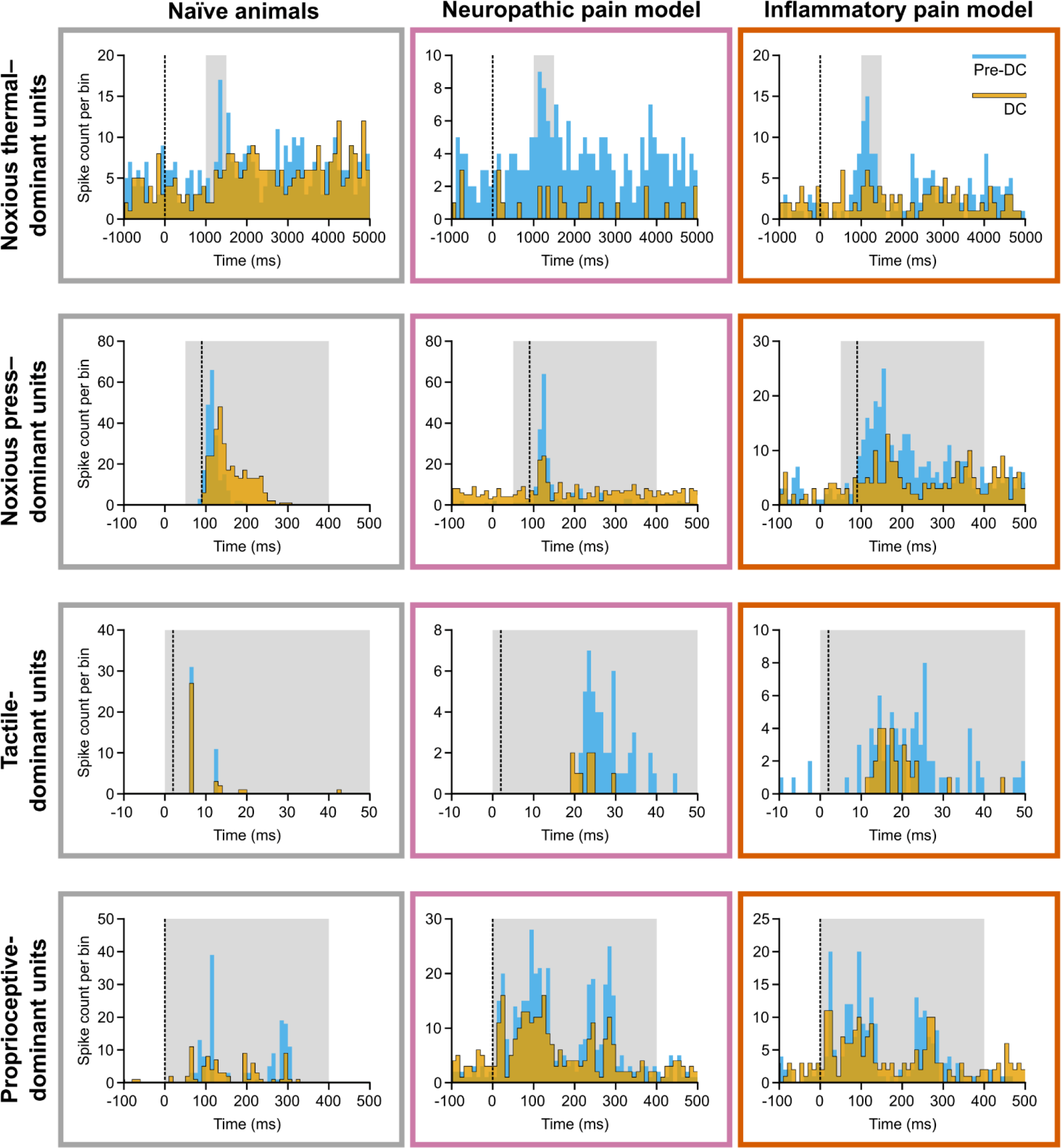
Example histograms of spike-sorted units. Histograms of recordings during 1000 μA DC (yellow) are overlaid on histograms of pre-DC baseline recordings (blue) of the same unit. Vertical dotted line indicates start of stimulus. Light grey box shows post-stimulus window in which spike events were totalled for analysis. Histogram bins are 1/60th of the displayed time (i.e. 100 ms for noxious thermal–dominant response, 10 ms for noxious press– and proprioceptive-dominant responses, and 1 ms for tactile-dominant response). Note that Y-axis is different for each plot to highlight within-panel differences rather than variance in binned peak.

**Table 1:**
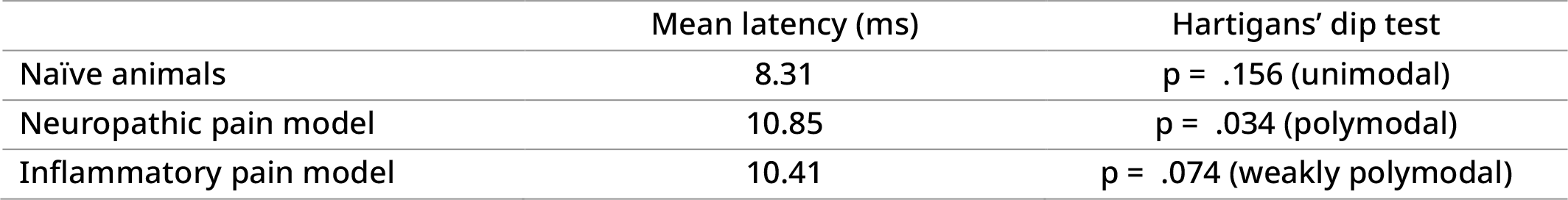
Distribution of response latencies for tactile-dominant units by treatment group. Interpretations of Hartigans’ dip test p-values are from Freeman and Dale (30).

In order to quantify the degree of DC-mediated suppression, we totalled spike events across the response window indicated by the grey box in each histogram of Figure 3. We analysed these windowed spike event totals using linear mixed-effects regressions, with significant differences determined by ANOVAs. Responses collected during application of DC at 500 μA and 1000 μA were compared against baseline responses prior to DC application (Figure 4). No significant differences in spike totals were found between cathodic and anodic DC; therefore, polarity was grouped for further analysis. We observed that at 500 μA DC, there was significant reduction in the post-stimulus activity of noxious thermal– (Figure 4A) and noxious press–dominant (Figure 4B) units in both pain model groups but not in naïve animals (p < .001 for pain models). At this amplitude, there was also reduction of tactile-dominant responses in the neuropathic pain group (Figure 4C, middle; p < .001) and proprioceptive-dominant responses in naïve animals (Figure 4D, left; p = .003). Additionally, at 1000 μA, there was significant suppression of noxious thermal–dominant responses in naïve animals (Figure 4A, left; p = .049) and tactile-dominant responses in the inflammatory pain group (Figure 4C, right; p = .008).

**Figure 4:**
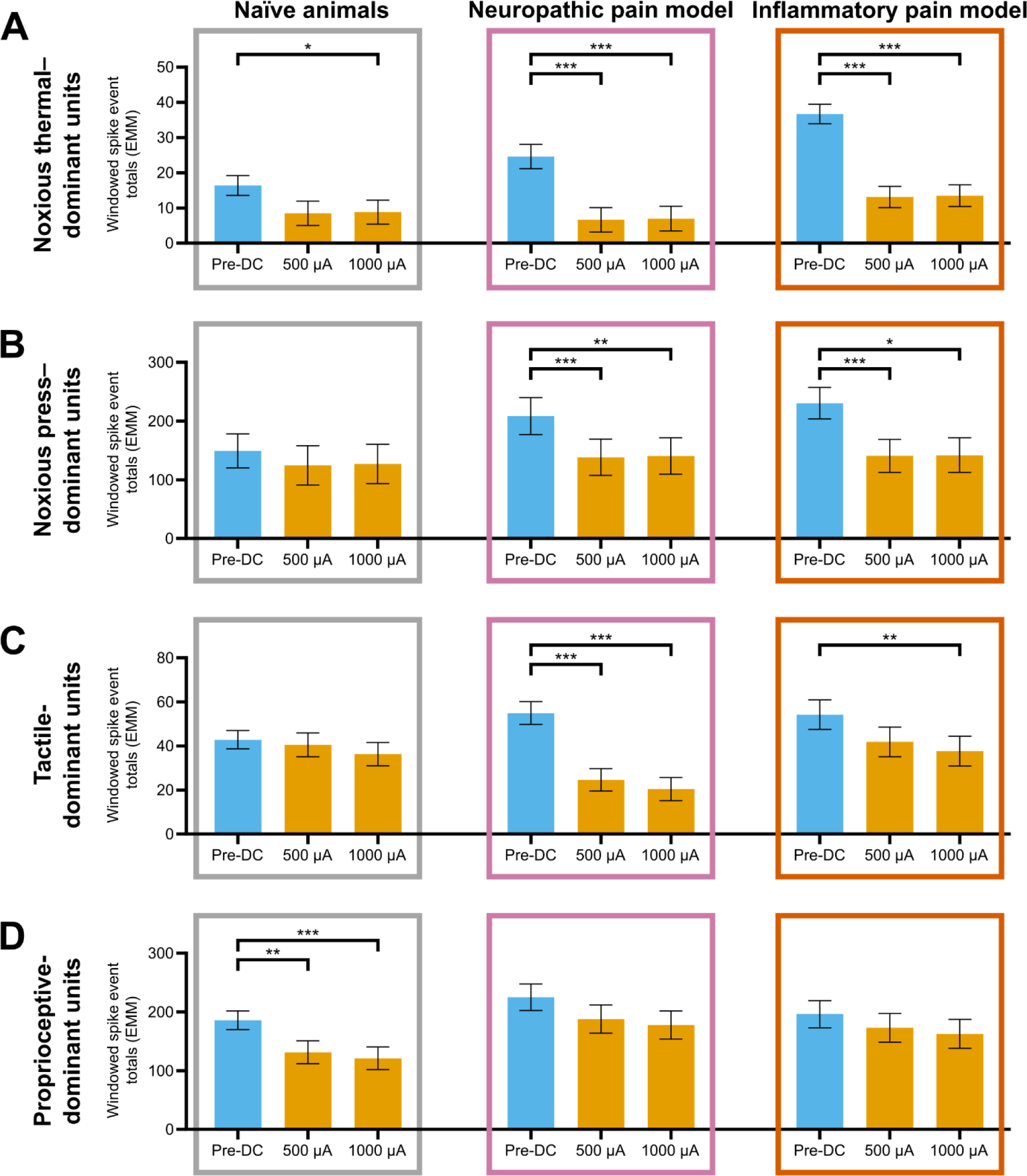
Windowed spike event totals before and during DC application. Spike totals were taken from post-stimulus windows, represented in Figure 3 by grey boxes. EMM ± SE of these totals are shown here, as derived from linear mixed-effects regression analyses. Each row A-D represents a different physiological stimulus modality, while each column corresponds to a treatment group. *: p < .05; **: p < .01; ***: p < .001. (A) Noxious thermal–dominant units. Significant differences were found between pre-DC and 1000 μA for all treatments, and between pre-DC and 500 μA in both pain model groups. Percentage reduction from baseline for these units in pain model groups were the largest of any test condition, at 63.1-73.1%. (B) Noxious press–dominant units. Significant differences were found between pre-DC and both DC amplitudes in pain models groups. No change from baseline was observed in naïve animals. (C) Tactile-dominant units. Significant differences were found between pre-DC and 1000 μA in both pain model groups, as well as between pre-DC and 500 μA in the neuropathic pain group. Reductions from baseline were relatively large in the neuropathic pain group, at 55.1-62.8%. (D) Proprioceptive-dominant units. Significant differences were found between pre-DC and both DC amplitudes in naïve animals, but not in either pain model group.

To compare the magnitude of DC-mediated suppression between test conditions, we calculated the percentage reduction from baseline wherever such a change was significant. The greatest reductions from baseline were seen in noxious thermal–dominant units in pain model treatment groups (63.1 – 73.1%) and tactile-dominant units in the neuropathic pain group (55.1 – 62.8%). Noxious thermal– dominant responses in naïve animals were reduced by 46.0% at 1000 μA, and noxious press–dominant responses in pain model groups were reduced by 32.6 – 38.9% during DC application. Tactile-dominant responses in the inflammatory pain group were reduced by 30.7% at 1000 μA and proprioceptive-dominant units in naïve animals were reduced by 29.0 – 34.4%.

In summary, DC neuromodulation significantly suppressed spinal responses evoked by different stimuli in naïve animals compared to pain model groups. The greatest reductions in activity were observed in noxious thermal–dominant units across all treatment groups as well as tactile-dominant units in the neuropathic pain group, with smaller reductions in noxious press–dominant units in both pain model groups and proprioceptive-dominant units in naïve animals.

### Pain-related spinal responses remain suppressed for longer than innocuous spinal responses following DC cessation

Previous studies found a sustained period of suppression after DC neuromodulation had ended. To examine this effect in our data, we first limited the dataset to units which were completely suppressed (defined as no significant difference between pre-stimulus and post-stimulus spike counts) during application of DC. A total of 36/38 noxious thermal–, 8/69 noxious press–, and 8/121 tactile-dominant units were identified from evoked units for which post-DC recordings were made. No proprioceptive-dominant units (0/87) were fully suppressed in our dataset. We observed recovery of activity in only 11 of the 36 suppressed noxious thermal–dominant units, while all noxious press– and tactile-dominant units recovered within 15 minutes. We compared the duration of post-DC block between these responses (Figure 5) and found that post-DC suppression persisted for longer in noxious thermal– (p < .001) and noxious pinch–dominant (p = .001) units than in tactile-dominant units (Mann-Whitney U).

**Figure 5:**
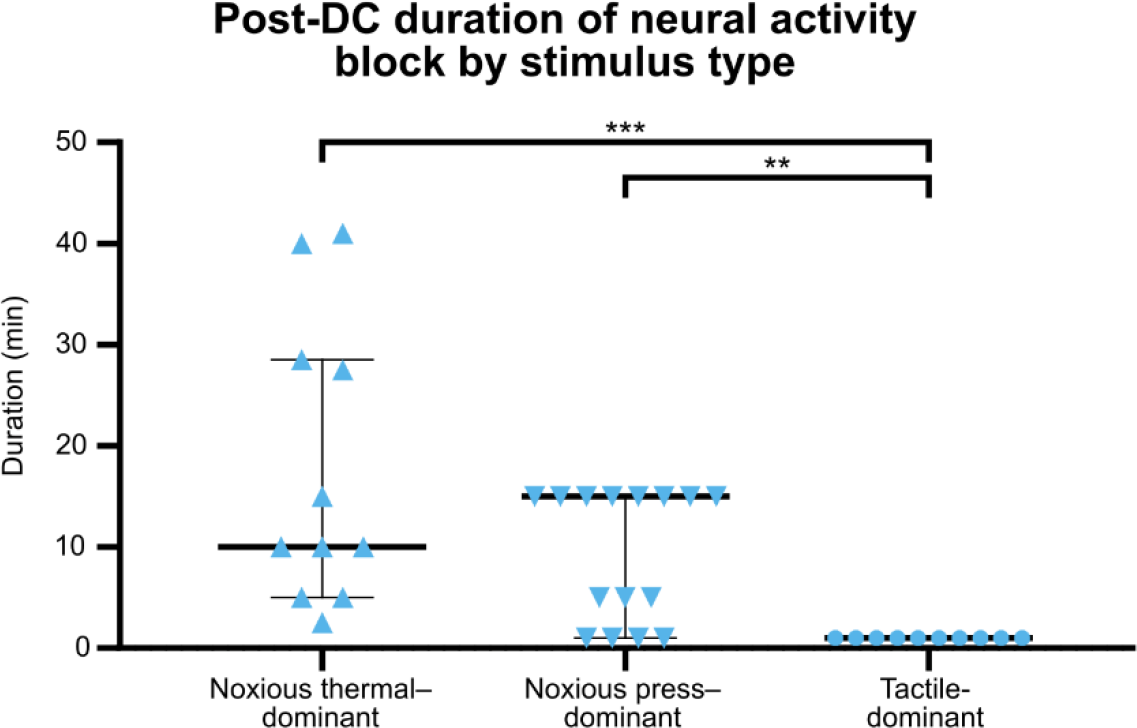
Post-DC duration of neural activity suppression. Time taken for evoked responses to reappear in units whose activity was completely suppressed during application of DC (p < .05). Proprioceptive-dominant units are not shown, as no such units were fully suppressed by DC. Significant differences in the distribution of post-DC neural activity suppression durations were found between tactile-dominant units and both noxious thermal– and noxious press– dominant units. **: p < .01; ***: p < .001.

## Discussion

The primary aim of this study was to compare the effects of peripheral DC neuromodulation on physiologically evoked spinal responses in rodent models of pain. We found that evoked responses relating to both hyperalgesia and allodynia were suppressed in pain models. Our results are broadly consistent with previous literature (18, 24) and provide evidence for the efficacy of peripheral DC in suppressing activity associated with chronic nociceptive and chronic neuropathic pain.

We found that noxious-evoked responses (thermal and high-threshold mechanical) were consistently suppressed in both pain models, and to a higher degree than in naïve animals. Noxious stimuli are associated with activity from small-diameter Aδ and C fibres, which are not typically sensitive to traditional pulsatile neuromodulation. Previous studies reported that DC could suppress these fibres at thresholds similar to those in large-diameter Aα and Aβ fibres (18, 24). Our results expand upon those findings in that we also demonstrate a functional suppression of nociceptive-associated activity at the spinal cord following peripheral DC neuromodulation. We achieved suppression at amplitudes roughly comparable to both Yang et al. (18) and Jones et al. (24). In nociceptive responses we observed near-maximal suppression at 500 µA, suggesting that lower amplitudes may also provide effective suppression. However, our study design did not explore peripheral mechanisms of action, and there may be contribution from indirect mechanisms such as desynchronization of spinal inputs or post-synaptic inhibition (31, 32).

Our data also showed increased responses to noxious stimuli in both pain models, which may reflect hyperalgesia associated with these pain states (33). There is some discrepancy between our electrophysiology and our behavioural verification as we did not find decreased thermal withdrawal thresholds in our neuropathic pain model. This is not unexpected, as thermal hyperalgesia does not consistently show in behavioural tests following spared tibial nerve injury, although the reason is not well understood (34–36). Our data nonetheless suggests that some hypersensitivity is present in these models at the dorsal horn, and that these responses are reduced by peripheral DC. These results are encouraging for the use of DC neuromodulation to suppress hyperalgesic responses in both neuropathic and nociceptive chronic pain states.

In contrast, there was a less straightforward effect in innocuous-evoked responses, with suppression of tactile-dominant activity in pain models, but not in naïve animals. Chronic pain, particularly chronic neuropathic pain, contains an allodynic component wherein previously innocuous stimuli are associated with painful sensations (26). Mechanisms of allodynia are complex, and involve pathological expression of low-threshold tactile receptors such as Piezo2 in high-threshold nociceptors as well as contribution of C-tactile afferents to painful sensation (37, 38). These mechanisms alter the function of small-diameter neurons and lead to decreased pain thresholds. We found significant changes in response patterns to innocuous tactile stimuli between our naïve animals and pain models, suggesting pathological changes in the tactile-evoked neural population. This is further supported by the increased baseline response seen in our pain model groups (Figure 4C), which displayed mechanical allodynia in behavioural tests. As such, we propose that peripheral DC may potentially suppress hypersensitive responses specifically associated with allodynia. However, further study is needed to confirm which populations of tactile-responding neurons are more strongly affected by peripheral DC.

We also observed suppression of proprioceptive-dominant responses in naïve animals, but the magnitude of suppression was smaller than for the other response types. We also did not find evidence of complete suppression in any proprioceptive unit. We expected some proprioceptive suppression to occur given similar findings by Jones et al. (24) but they observed this effect in neuropathic pain models, and with a complete block, while we did not. These results reinforce that DC neuromodulation is sensitive to small changes such as stimulation site and electrode position (32, 39). Regardless, we recommend that any future chronic stimulation study incorporates gait analysis to determine whether any potential proprioceptive suppression results in functional deficit.

Both Yang et al. (18) and Jones et al. (24) reported long C fibre recovery times following DC block. Jones et al. utilised this phenomenon to develop an alternative method for selective block by delivering ultra-low frequency waveforms which allow only innocuous signalling to recover between phases. In the present study we show comparable results, with some thermal-dominant responses only returning 30 minutes or more after DC delivery was stopped. This provides further evidence to support their approach. However, it reveals a potential confound in our study design. Although we show long time courses for recovery, sequential recordings in multiple locations are made over the course of an experiment. As the DC waveform was delivered to the entire sciatic nerve, it is possible that its effects persisted when starting a new set of recordings. We controlled for this by leaving long recovery periods and always comparing against pre-DC baseline activity. We cannot completely rule out the possibility of longer duration effects of DC neuromodulation, but their analysis is beyond the scope of this study.

By demonstrating suppression of functional, pain-related activity in both neuropathic and inflammatory nociceptive pain models, we highlight an advantage of peripheral DC neuromodulation over traditional pulsatile stimulation. There is limited evidence for the efficacy of existing devices in the management of chronic nociceptive pain conditions (13, 14), such as post-surgical or arthritic pain. DC neuromodulation could therefore be particularly valuable for the treatment of these conditions, and help to minimise risks associated with chronic opioid use (5, 40). To this end, our next steps include development of a chronically implantable device with which we will test long-term implantable lead safety. In parallel, we will perform awake behaving animal experiments to further validate functional suppression of nociception and pain. Overall, our results support the continued translation of peripheral DC neuromodulation as a potential treatment for chronic pain.

## Materials and methods

### Animals

Experiments were performed on Sprague Dawley rats aged 10-23 weeks of both sexes (25 females, 235-387 g; 22 males, 453-720 g) obtained from the Animal Resources Centre (WA, AUS). Rats were housed on a 12-hour light-dark cycle with ad libitum access to food and water, in groups of up to four according to sex. All procedures were approved by the University of New South Wales Animal Care and Ethics Committee (20/80A).

### Pain models

#### Neuropathic pain model

A neuropathic pain state was induced using the spared nerve injury (SNI) model described by Decosterd and Woolf (41), with the tibial branch of the sciatic nerve spared instead of the sural branch (35, 36). Sparing of the tibial branch maintained hindpaw sensation across a larger dermatome whilst preserving neuropathic pain behaviours resulting from the injury. Rats were anesthetised with isoflurane (4% induction, 1.5% maintenance) in oxygen (2 L/min induction, 1 L/min maintenance). The common peroneal and sural nerves on the left side were ligated with Mersilk 5/0 suture (Ethicon; NJ, USA) and axotomised distally, while the tibial nerve was left intact. The wound was sealed and antiseptic cream applied. Rats were monitored until awake and behaving normally. General health and wellbeing were monitored daily following the procedure until the terminal electrophysiology experiments 8 days later, when peak pain hypersensitivity was achieved (35, 41).

#### Inflammatory pain model

An inflammatory pain state was induced by subcutaneous injection of complete Freund’s adjuvant (CFA; 42). Rats were anesthetised as above and 100 μL CFA (Sigma-Aldrich; MA, USA) was injected subcutaneously into the plantar surface of the left hindpaw to produce a local inflammatory reaction. Post-operative pain monitoring was performed as per neuropathic pain models, with the exception that the terminal experiment was instead conducted 5 days after CFA injection to correspond to peak pain hypersensitivity in this model (43).

### Behavioural tests

Sensitivity to mechanical allodynia and thermal hyperalgesia were tested before injury to establish baseline, and again on day 7 for SNI-treated rats, or day 4 for CFA-treated rats; different time courses were chosen to allow peak pain hypersensitivity to develop in each injury (35, 41, 43). Testing was performed after 15 minutes of habituation and in well-controlled environments to minimise non-specific responses (44). Withdrawal reflexes were defined as sudden lifting and/or licking of the hindpaw in both tests.

To verify the development of mechanical allodynia, rats were habituated in an enclosure on an elevated mesh grid. Withdrawal threshold was measured by pressing an electronic von Frey aesthesiometer (IITC Life Science Inc.; CA, USA) into the plantar hindpaw until a withdrawal reflex was evoked (44); the maximal force applied was recorded. Three such measurements were taken with 2 minutes between trials.

To verify the development of thermal hyperalgesia, rats were instead habituated in a glass-bottomed cage. A 50 mW/cm2 infrared heat source (Heat-Flux Radiometer; Ugo Basile; VA, ITA) was applied to the plantar surface of the hindpaw until a withdrawal reflex was evoked and the withdrawal latency recorded (45). A pre-determined cut-off latency of 30 seconds was used to prevent tissue damage. Three trials were performed with 2 minutes between trials.

### Electrophysiology

Rats were anesthetised using urethane (1.5 g/kg body weight; Sigma-Aldrich; MA, USA) administered via intraperitoneal injection. The animal was placed on a heating block (36.5 °C) and 10 mL/kg body weight of modified Krebs-Henseleit solution (117.9 mM NaCl, 4.7 mM KCl, 25 mM NaHCO3, 1.3 mM NaH2PO4, 1.2 mM MgSO4, 2.5 mM CaCl2) was injected subcutaneously every 3 hours to maintain hydration. The animal was intubated by tracheotomy, fixed to a stereotaxic frame (Stoelting; IL, USA) and a laminectomy performed to expose the T13 and L1 regions of the dorsal spinal cord. The dura mater and arachnoid mater covering the exposed spinal cord were removed, and a 32-channel penetrating multielectrode array (NeuroNexus; MI, USA) was inserted approximately 1 mm deep using a micromanipulator ipsilateral to the stimulation site (Figure 1D-F). Voltage signals were band-pass filtered from 0.1 Hz to 20 kHz by an RHS Stim/Recording Headstage and recorded at 30 kHz using an RHS Stim-Recording System (Intan Technologies; CA, USA). To visualise the depth of electrode penetration, the MEA was soaked in DiI (Sigma-Aldrich; MA, USA) before insertion into the spinal cord in three animals (Figure 1F). At the end of the experiment, those animals were perfused via 4% paraformaldehyde and their spinal cords removed, cryoprotected, and sectioned at 10 µm on a Leica CM1950 cryostat (HE, DEU). Sections were imaged using an Olympus IX83 Inverted Microscope (MA, USA).

### Peripheral stimulation

#### Noxious thermal–dominant stimulus

To evoke a thermoreceptive-dominant response, A 980 nm continuous wave diode laser (Changchun New Industries Optoelectronics Technology Co., Ltd.; JL, CHN) with a beam diameter of 4 mm was positioned normal to the plantar surface of the hindpaw. A 1 second beam was given at 1.95-6.85 W to heat the stimulation site by ∼10 °C, as verified by a ThermaCAM Reporter7 Pro thermal camera (FLIR Systems; OR, USA). Sets of two laser pulses were recorded, with the inter-pulse interval ranging from 20 seconds to 5 minutes.

#### Noxious press–dominant stimulus

To evoke a high-threshold (noxious) mechanoreceptive-dominant response, a custom piezoelectric pressure sensor was fixed to the table and the hindpaw placed upon it with the plantar surface facing upwards. A wooden rod was used to firmly press the stimulation site at a rate of ∼1 Hz. 5 sets of 10 repetitions were recorded alongside analogue voltage changes at the piezoelectric sensor (46).

#### Tactile-dominant stimulus

To evoke a low threshold (innocuous) mechanoreceptive-dominant response a lightweight aluminium rod with a rounded plastic cap was fixed to a SignalForce V4 shaker (Data Physics; CA, USA) and positioned with its tip resting on the plantar surface of the hindpaw. Mechanical stimulation waveforms were passed through a SignalForce PA100E linear amplifier (Data Physics; CA, USA) to the shaker, which was calibrated for a peak downwards displacement of 100 μm over 250 μs using a laser displacement sensor (Micro-Epsilon Messtechnik; BY, DEU). Mechanical stimulation was given at a rate of 1 Hz in 5 sets of 10 repetitions.

#### Proprioceptive-dominant stimulus

To evoke a proprioceptive-dominant response, a wooden rod attached to a stepper motor (Pololu Corporation; NV, USA) was placed under the hindpaw, supporting the ankle. The motor rotated 15° over a 200 ms period before returning to its initial position, causing bilateral rotation of the ankle and knee joints. 5 sets of 10 rotations at 1 Hz were delivered.

### DC neuromodulation

The ipsilateral biceps femoris and gluteus superficialis were separated to expose a section of the sciatic nerve ipsilateral to the stimulation site (47). A silicone cuff consisting of three electrolytic agar gel channels (a central active channel and two flanking returns) was inserted around the sciatic nerve. A tripolar cuff design was selected to minimise onset activation (48). These channels were connected to a Model 2100 or 4100 isolated stimulator (A-M Systems; WA, USA) using stainless steel leads suspended in saline. A DC waveform with 10s on-/offset ramp was delivered through the cuff (Figure 1H). Two amplitudes were used (500 μA and 1000 μA) for both cathodic (−) and anodic (+) active channel polarities. The total duration of the waveform was 100-110 seconds, but a duration of up to 5 minutes was used for noxious thermal units in naïve animals. There were three experimental phases in relation to DC application: pre-DC recordings to establish baseline activity, during-DC recordings to determine level of suppression, and post-DC recovery recordings to observe return to baseline. MRI images were collected from a separate cohort of two animals that were implanted with the tripolar silicone cuff and allowed to recover over the course of 1 week (Figure 1G). Images were taken at the Biological Resources Imaging Laboratory (UNSW; NSW, AUS) under isoflurane anaesthesia.

### Spike-sorting and cell classification

Signal processing of electrophysiological recordings was performed using custom Julia scripts (v1.8; 49). Signals were band-pass filtered from 300-5000 Hz and separated into trials. Sets of three spatially adjacent channels were extracted for further analysis based on the presence and volume of post-stimulus spike activity (7 additional spike events and at least twice the number of spike events compared to pre-DC baseline). A pre-DC recording from each set was designated as the template and the principal components of its concatenated waveforms clustered using an unsupervised k-means algorithm (50).

When 1% or more inter-spike intervals for a given cluster were ≤1 ms, it was isolated and reiterated through the k-means algorithm; this was performed up to four times per cluster before exclusion. The final clustering solution was then applied to other sets from the same response. Units were classified according to the stimulus paradigm delivered in the template set. Noxious press– and proprioceptive-dominant stimuli unavoidably contained a tactile component and sometimes evoked low-threshold tactile responses. To avoid misclassification, non–tactile-dominant units were discarded if they also showed a significant response to tactile-dominant stimulus. Spike events in pre- and post-stimulation windows were counted and exported for statistical analysis; window timings varied according to stimulation paradigm (Table 2).

**Table 2:**
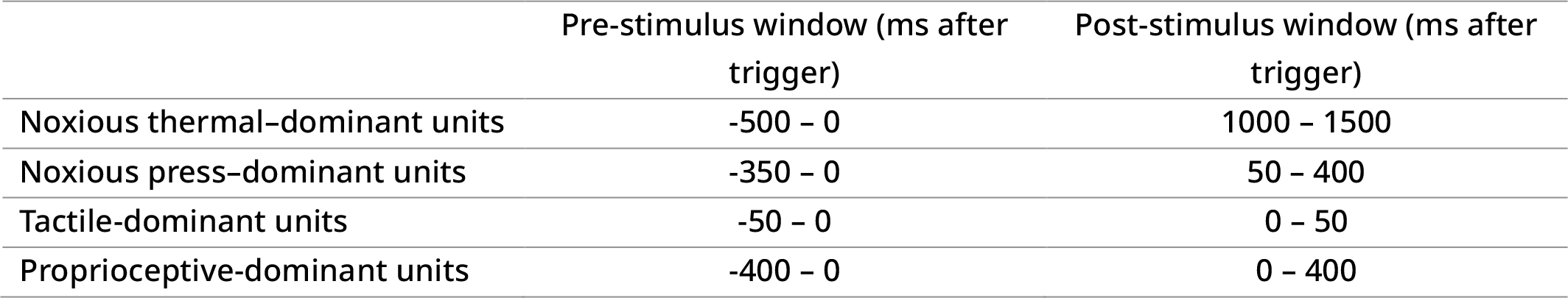
Window timings for analysis of post-stimulus spike-event totals.

### Statistical analysis

Custom R scripts (v4.2.2) were used for statistical analysis. For behaviour, measurements were analysed using linear mixed-effects regression with Satterthwaite’s method (LMER; lmer function from the lmerTest package in R) and analysis of variance (ANOVA) to determine significant differences (α = .05). Fixed effects included experimental phase, treatment group, and sex. Animal identifiers were used as a random effect.

For electrophysiological recordings, analysis was performed using the same LMER package, with a separate model for each stimulation paradigm. Fixed effects included experimental phase, treatment group, presence of response (pre-vs. post-stimulus events, p < .05), sex, DC amplitude, time since DC, and pre-stimulus spike count. Animal and unit identifiers were used as random effects. Model fitting was assessed by residual plot inspection. Where significant effects were found, multiple comparisons analysis using Tukey p-value adjustments were made (emmeans function from the emmeans package in R). Data are visualised as estimated marginal means with standard error (EMM ± SE). Effects of polarity were analysed using a separate LMER, with DC amplitude, treatment group, presence of response, sex, time since DC, and pre-stimulus spike count as fixed effects, and animal and unit identifiers as random effects. Latency of the response peak was analysed in tactile-dominant units using Kruskal-Wallis and Hartigans’ dip tests. A Wilcoxon rank-sum test analysis was also performed on units fully suppressed by DC (defined as no significant difference between pre-stimulus and post-stimulus spike counts) to compare the duration after DC cessation for which complete suppression persisted. All significance was defined as p < .05.

## Data availability

Data files and code have been made available at the following Git repository: https://github.com/Aplinlab/Su-et-al.-2023-Data

## Funding

National Health and Medical Research Council of Australia Ideas Grant (APP1187416) Felix P. Aplin. The funders had no role in study design, data collection and interpretation, or the decision to submit the work for publication.

## Acknowledgements

We would like to acknowledge C. Cheng and P. Adkisson for their assistance with device design and fabrication. We would also like to thank I. Birznieks, P. Carrive, and R. Vickery for the generous loan of their equipment, as well as A. Fernando and Z. Zheng for assistance with coding.

---

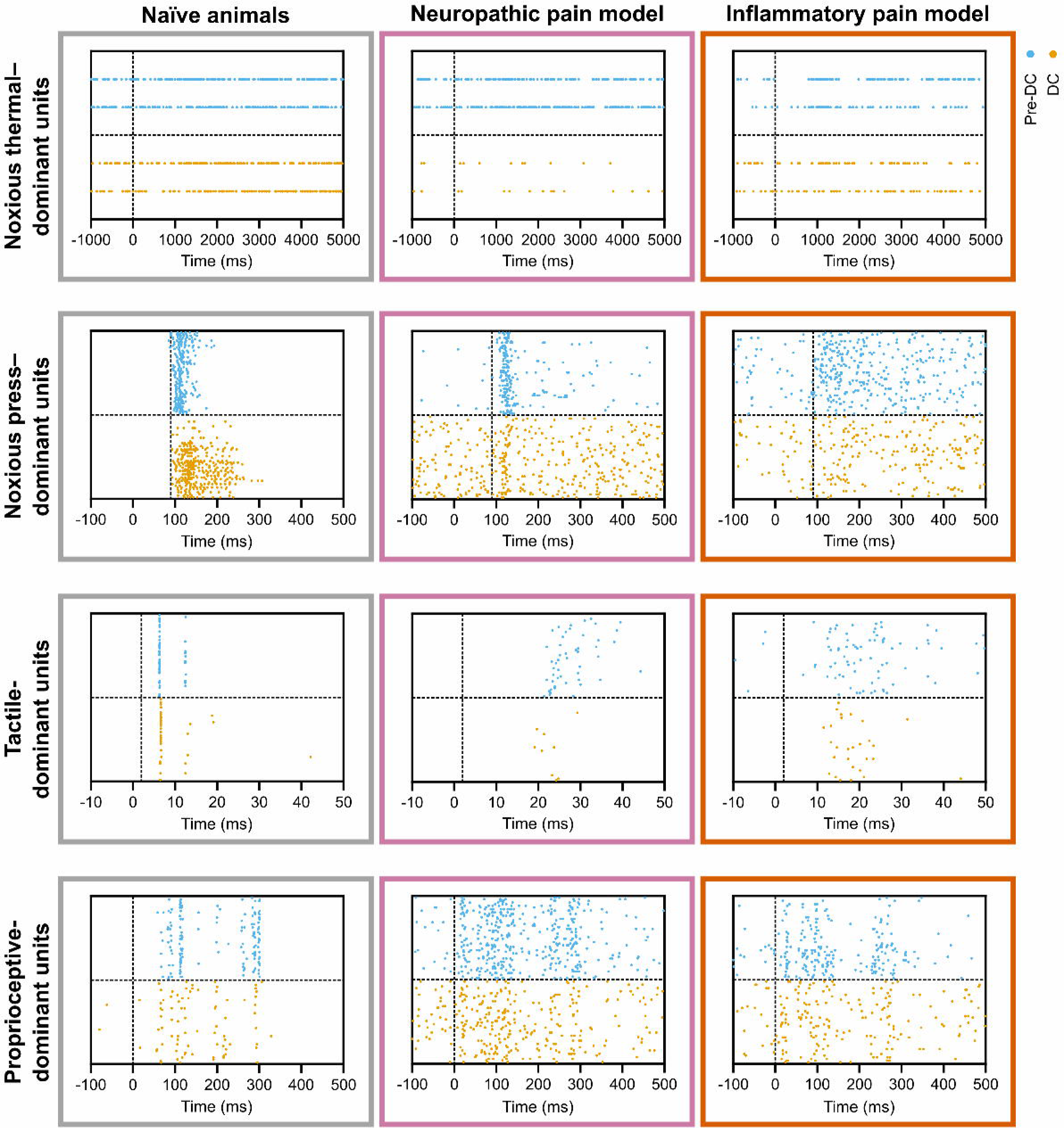

## Notes

https://github.com/Aplinlab/Su-et-al.-2023-Data

### Competing Interest Statement

MNS, GMT, GYF, and FPA have filed a provisional patent application related to this paper ("Electrode Lead and method for pain blocking using ionic direct current"; US63,493,50; Provisional.). All other authors declare they have no competing interests.

### Summary of Updates

Reformatted and restructured manuscript for submission to eLife. No major changes to the text other than a rewritten abstract.

## References

1. R. J. Yong, P. M. Mullins, N. Bhattacharyya, Prevalence of chronic pain among adults in the United States. Pain 163, e328–e332 (2022).

2. R. D. Treede, W. Rief, A. Barke, Q. Aziz, M. I. Bennett, R. Benoliel, M. Cohen, S. Evers, N. B. Finnerup, M. B. First, M. A. Giamberardino, S. Kaasa, B. Korwisi, E. Kosek, P. Lavand’homme, M. Nicholas, S. Perrot, J. Scholz, S. Schug, B. H. Smith, P. Svensson, J. W. S. Vlaeyen, S.-J. Wang, Chronic pain as a symptom or a disease: the IASP Classification of Chronic Pain for the International Classification of Diseases (ICD-11). PAIN 160, 19–27 (2019).

3. R. D. Treede, W. Rief, A. Barke, Q. Aziz, M. I. Bennett, R. Benoliel, M. Cohen, S. Evers, N. B. Finnerup, M. B. First, M. A. Giamberardino, S. Kaasa, E. Kosek, P. Lavand’homme, M. Nicholas, S. Perrot, J. Scholz, S. Schug, B. H. Smith, P. Svensson, J. W. S. Vlaeyen, S. J. Wang, A classification of chronic pain for ICD-11. Pain 156, 1003–1007 (2015).

4. N. Hylands-White, R. V. Duarte, J. H. Raphael, An overview of treatment approaches for chronic pain management. Rheumatol Int 37, 29–42 (2017).

5. R. Ward, D. Taber, H. Gonzales, M. Gebregziabher, W. Basco, J. McCauley, P. Mauldin, S. Ball, Risk factors and trajectories of opioid use following total knee replacement. Knee Surg Relat Res 34, 18 (2022).

6. M. Kimura, J. C. Eisenach, K. I. Hayashida, Gabapentin loses efficacy over time after nerve injury in rats: role of glutamate transporter-1 in the locus coeruleus. Pain 157, 2024–2032 (2016).

7. T. R. Deer, M. F. Esposito, W. P. McRoberts, J. S. Grider, D. Sayed, P. Verrills, T. J. Lamer, C. W. Hunter, K. V. Slavin, J. M. Shah, J. M. Hagedorn, T. Simopoulos, D. A. Gonzalez, K. Amirdelfan, S. Jain, A. Yang, R. Aiyer, A. Antony, N. Azeem, R. M. Levy, N. Mekhail, A Systematic Literature Review of Peripheral Nerve Stimulation Therapies for the Treatment of Pain. Pain Med 21, 1590–1603 (2020).

8. A. D. Kaye, S. Ridgell, E. S. Alpaugh, A. Mouhaffel, A. J. Kaye, E. M. Cornett, A. A. Chami, R. Shah, B. M. Dixon, O. Viswanath, I. Urits, A. N. Edinoff, R. D. Urman, Peripheral Nerve Stimulation: A Review of Techniques and Clinical Efficacy. Pain Ther 10, 961–972 (2021).

9. K. Kumar, R. S. Taylor, L. Jacques, S. Eldabe, M. Meglio, J. Molet, S. Thomson, J. O’Callaghan, E. Eisenberg, G. Milbouw, E. Buchser, G. Fortini, J. Richardson, R. B. North, Spinal cord stimulation versus conventional medical management for neuropathic pain: a multicentre randomised controlled trial in patients with failed back surgery syndrome. Pain 132, 179–188 (2007).

10. S. F. Lempka, P. G. Patil, Innovations in spinal cord stimulation for pain. Current Opinion in Biomedical Engineering 8, 51–60 (2018).

11. M. Hofmeister, A. Memedovich, S. Brown, M. Saini, L. E. Dowsett, D. L. Lorenzetti, T. L. McCarron, G. MacKean, F. Clement, Effectiveness of Neurostimulation Technologies for the Management of Chronic Pain: A Systematic Review. Neuromodulation 23, 150–157 (2020).

12. H. Knotkova, C. Hamani, E. Sivanesan, M. F. E. Le Beuffe, J. Y. Moon, S. P. Cohen, M. A. Huntoon, Neuromodulation for chronic pain. Lancet 397, 2111–2124 (2021).

13. B. T. Sitzman, D. A. Provenzano, Best Practices in Spinal Cord Stimulation. Spine (Phila Pa 1976) 42 Suppl 14, S67–S71 (2017).

14. C. M. Stewart, M. Y. J. Qadri, C. A. Daly, Upper-Extremity Peripheral Nerve Stimulators. J Hand Surg Glob Online 5, 121–125 (2023).

15. J. Holsheimer, Which Neuronal Elements are Activated Directly by Spinal Cord Stimulation. Neuromodulation 5, 25–31 (2002).

16. N. H. Strand, R. D’Souza, C. Wie, S. Covington, M. Maita, J. Freeman, J. Maloney, Mechanism of Action of Peripheral Nerve Stimulation. Curr Pain Headache Rep 25, 47 (2021).

17. E. Pena, N. A. Pelot, W. M. Grill, Non-monotonic kilohertz frequency neural block thresholds arise from amplitude-and frequency-dependent charge imbalance. Sci Rep 11, 5077 (2021).

18. F. Yang, M. Anderson, S. He, K. Stephens, Y. Zheng, Z. Chen, S. N. Raja, F. Aplin, Y. Guan, G. Fridman, Differential expression of voltage-gated sodium channels in afferent neurons renders selective neural block by ionic direct current. Sci Adv 4, eaaq1438 (2018).

19. J. R. Bartlett, R. W. Doty, B. B. Lee, N. Negrao, W. H. Overman, Jr., Deleterious effects of prolonged electrical excitation of striate cortex in macaques. Brain Behav Evol 14, 46–66 (1977).

20. S. B. Brummer, J. McHardy, M. J. Turner, Electrical stimulation with Pt electrodes: Trace analysis for dissolved platinum and other dissolved electrochemical products. Brain Behav Evol 14, 10–22 (1977).

21. G. E. Foxworthy, G. Y. Fridman, in 2022 IEEE Biomedical Circuits and Systems Conference (BioCAS). (2022), pp. 321-325.

22. G. Y. Fridman, C. C. D. Santina, Safe Direct Current Stimulation to Expand Capabilities of Neural Prostheses. IEEE Transactions on Neural Systems and Rehabilitation Engineering 21, 319–328 (2013).

23. T. Vrabec, N. Bhadra, G. Van Acker, N. Bhadra, K. Kilgore, Continuous Direct Current Nerve Block Using Multi Contact High Capacitance Electrodes. IEEE Trans Neural Syst Rehabil Eng 25, 517–529 (2017).

24. M. G. Jones, E. R. Rogers, J. P. Harris, A. Sullivan, D. M. Ackermann, M. Russo, S. F. Lempka, S. B. McMahon, Neuromodulation using ultra low frequency current waveform reversibly blocks axonal conduction and chronic pain. Sci Transl Med 13, (2021).

25. F. P. Aplin, G. Y. Fridman, Implantable Direct Current Neural Modulation: Theory, Feasibility, and Efficacy. Front Neurosci 13, 379 (2019).

26. J. N. Campbell, R. A. Meyer, Mechanisms of neuropathic pain. Neuron 52, 77–92 (2006).

27. H. Ueda, Peripheral mechanisms of neuropathic pain - involvement of lysophosphatidic acid receptor-mediated demyelination. Mol Pain 4, 11 (2008).

28. J. Scholz, Mechanisms of chronic pain. Mol Pain 10, O15 (2014).

29. J. A. Hartigan, P. M. Hartigan, The Dip Test of Unimodality. The Annals of Statistics 13, 70–84, 15 (1985).

30. J. B. Freeman, R. Dale, Assessing bimodality to detect the presence of a dual cognitive process. Behav Res Methods 45, 83–97 (2013).

31. T. Radman, Y. Su, J. H. An, L. C. Parra, M. Bikson, Spike Timing Amplifies the Effect of Electric Fields on Neurons: Implications for Endogenous Field Effects. The Journal of Neuroscience 27, 3030–3036 (2007).

32. D. Chakraborty, D. Q. Truong, M. Bikson, H. Kaphzan, Neuromodulation of Axon Terminals. Cereb Cortex 28, 2786–2794 (2018).

33. T. S. Jensen, N. B. Finnerup, Allodynia and hyperalgesia in neuropathic pain: clinical manifestations and mechanisms. Lancet Neurol 13, 924–935 (2014).

34. J. Fang, J. Du, X. Xiang, X. Shao, X. He, Y. Jiang, B. Liu, Y. Liang, J. Fang, SNI and CFA induce similar changes in TRPV1 and P2X3 expressions in the acute phase but not in the chronic phase of pain. Exp Brain Res 239, 983–995 (2021).

35. F. Guida, D. De Gregorio, E. Palazzo, F. Ricciardi, S. Boccella, C. Belardo, M. Iannotta, R. Infantino, F. Formato, I. Marabese, L. Luongo, V. de Novellis, S. Maione, Behavioral, Biochemical and Electrophysiological Changes in Spared Nerve Injury Model of Neuropathic Pain. Int J Mol Sci 21, (2020).

36. M. H. Ko, M. L. Yang, S. C. Youn, C. T. Lan, T. J. Tseng, Intact subepidermal nerve fibers mediate mechanical hypersensitivity via the activation of protein kinase C gamma in spared nerve injury. Mol Pain 12, (2016).

37. S. E. Murthy, M. C. Loud, I. Daou, K. L. Marshall, F. Schwaller, J. Kuhnemund, A. G. Francisco, W. T. Keenan, A. E. Dubin, G. R. Lewin, A. Patapoutian, The mechanosensitive ion channel Piezo2 mediates sensitivity to mechanical pain in mice. Sci Transl Med 10, (2018).

38. S. S. Nagi, T. K. Rubin, D. K. Chelvanayagam, V. G. Macefield, D. A. Mahns, Allodynia mediated by C-tactile afferents in human hairy skin. J Physiol 589, 4065–4075 (2011).

39. M. Bikson, M. Inoue, H. Akiyama, J. K. Deans, J. E. Fox, H. Miyakawa, J. G. Jefferys, Effects of uniform extracellular DC electric fields on excitability in rat hippocampal slices in vitro. J Physiol 557, 175–190 (2004).

40. J. A. Zamora-Legoff, S. J. Achenbach, C. S. Crowson, M. L. Krause, J. M. Davis3rd, E. L. Matteson, Opioid use in patients with rheumatoid arthritis 2005-2014: a population-based comparative study. Clin Rheumatol 35, 1137-1144 (2016).

41. I. Decosterd, C. J. Woolf, Spared nerve injury: an animal model of persistent peripheral neuropathic pain. Pain 87, 149–158 (2000).

42. E. Claassen, W. de Leeuw, P. de Greeve, C. Hendriksen, W. Boersma, Freund’s complete adjuvant: an effective but disagreeable formula. Res Immunol 143, 478–483; discussion 572 (1992).

43. E. Eliav, R. Benoliel, M. Tal, Inflammation with no axonal damage of the rat saphenous nerve trunk induces ectopic discharge and mechanosensitivity in myelinated axons. Neurosci Lett 311, 49–52 (2001).

44. J. R. Deuis, L. S. Dvorakova, I. Vetter, Methods Used to Evaluate Pain Behaviors in Rodents. Front Mol Neurosci 10, 284 (2017).

45. K. Hargreaves, R. Dubner, F. Brown, C. Flores, J. Joris, A new and sensitive method for measuring thermal nociception in cutaneous hyperalgesia. Pain 32, 77–88 (1988).

46. A. J. Loutit, J. R. Potas, Dorsal Column Nuclei Neural Signal Features Permit Robust Machine-Learning of Natural Tactile- and Proprioception-Dominated Stimuli. Frontiers in Systems Neuroscience 14, (2020).

47. P. J. Austin, A. Wu, G. Moalem-Taylor, Chronic Constriction of the Sciatic Nerve and Pain Hypersensitivity Testing in Rats. JoVE, e3393 (2012).

48. E. Pena, N. A. Pelot, W. M. Grill, Spatiotemporal parameters for energy efficient kilohertz-frequency nerve block with low onset response. J Neuroeng Rehabil 20, 72 (2023).

49. J. Bezanson, A. Edelman, S. Karpinski, V. B. Shah, Julia: A fresh approach to numerical computing. SIAM review 59, 65–98 (2017).

50. J. R. Potas, N. G. de Castro, T. Maddess, M. N. de Souza, Waveform Similarity Analysis: A Simple Template Comparing Approach for Detecting and Quantifying Noisy Evoked Compound Action Potentials. PLoS One 10, e0136992 (2015).

