## Supplementary Figure S1 for "Peripheral direct current suppresses physiologically evoked nociceptive responses at the spinal cord in rodent models of pain"

School of Biomedical Sciences, University of New South Wales, Sydney, Australia; Eccles Institute, John Curtin School of Medical Research, The Australian National University, Canberra, Australia; Graduate School of Biomedical Engineering, University of New South Wales, Sydney, Australia; Department of Otolaryngology, Head and Neck Surgery, Johns Hopkins University, Baltimore, United States; Department of Biomedical Engineering, Johns Hopkins University, Baltimore, United States; Department of Electrical and Computer Engineering, Johns Hopkins University, Baltimore, United States

#### Supplementary materials

This file includes:

- Supplementary Figure S1

### Figures

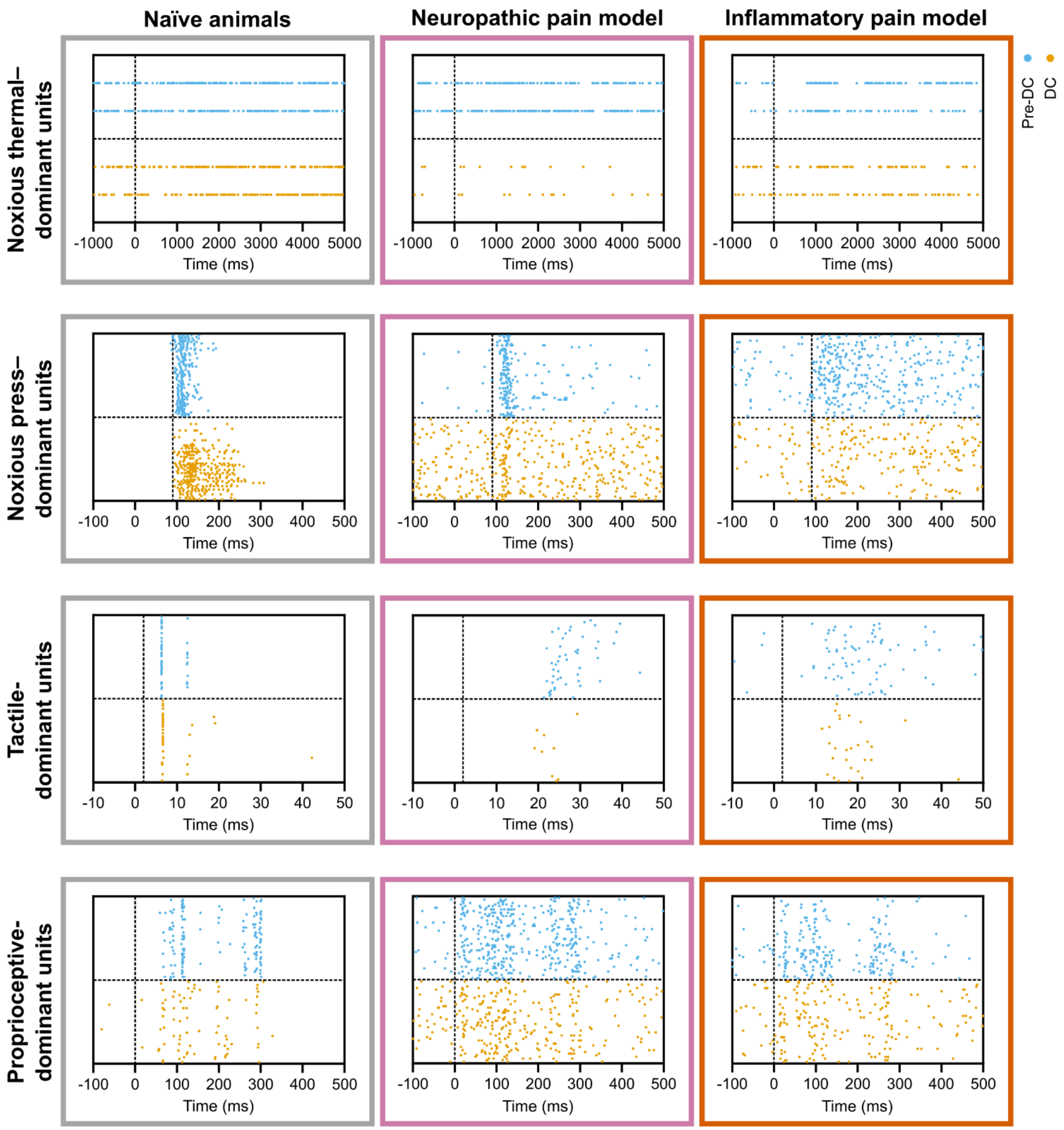


Figure S1: Example rasters of spike-sorted units.

Rasters of pre-DC baseline recordings (blue) and recordings during 1000 µA DC (yellow). Baseline–DC pairs are taken from the same unit. Each unit was recorded from a different animal. Vertical dotted line indicates start of stimulus.
